# Proteome-wide identification of amino acid substitutions deleterious for protein function

**DOI:** 10.1101/2022.04.06.487405

**Authors:** Ricard A. Rodriguez-Mias, Kyle N. Hess, Bianca Y. Ruiz, Ian R. Smith, Anthony S. Barente, Stephanie M. Zimmerman, Yang Y. Lu, William S. Noble, Stanley Fields, Judit Villén

**Affiliations:** Department of Genome Sciences, University of Washington, Graduate Program in Molecular and Cellular Biology, University of Washington

## Abstract

DNA sequencing has led to the discovery of millions of mutations that change the encoded protein sequences, but the impact of nearly all of these mutations on protein function is unknown. We addressed this scarcity of functional data by developing Miro, a proteomic technology that uses mistranslation to introduce amino acid substitutions and biochemical assays to quantify functional differences of thousands of protein variants by mass spectrometry. We apply this technology to the proteome of yeast to reveal amino acid substitutions that impact protein structure, ligand binding, protein-protein interactions, protein post-translational modifications, and protein thermal stability. Adapting Miro to human cells will provide a means to efficiently accelerate our mechanistic interpretation of genomic mutations to predict disease risk.

## Introduction

Changes to protein sequences as a consequence of mutations in DNA can alter protein structure and function, impact cellular and organismal physiology, and result in disease^1^. Thus, deciphering which amino acid substitutions have functional consequences is a major focus of studies in biology, protein engineering, and medical genetics. The predominant approach to determine the functional effect of a change to a protein sequence has been mutagenesis followed by measurement of the resulting mutant protein’s function. However, developments in genomics have accelerated the discovery of mutations such that tens of millions of missense variants in the human genome need functional interpretation^2–4^. Traditional mutagenesis approaches are severely underpowered for this task, and even recent high-throughput implementations, such as deep mutational scanning^5,6^, assess the effect of mutations in only one protein per experiment. Considering the ~20,000 human genes and the vast proteome encoded by these genes, such mutational experiments would require an inordinate amount of time to characterize the missense variants discovered to date.

To overcome this major bottleneck in genome analysis, we developed Miro, an approach to introduce amino acid substitutions into proteins and assess their consequences on protein function at a proteome-wide scale. First, Miro leverages the permissive nature of aminoacyl tRNA synthetases, which can incorporate closely related non-canonical amino acids (ncAAs) during protein translation in living systems ranging from bacteria to human cells^7–9^. Cells grown in the presence of ncAAs generate statistical proteomes composed of thousands of protein quasispecies, each defined as a collection of protein variants sharing most of their sequence, with the exception of the randomly introduced ncAAs. Although similar to their cognate counterparts, ncAAs can introduce significant changes to hydrophobicity, pKa, and secondary structure and thereby serve as a useful tool to probe for positional sensitivity in protein sequences. Second, Miro applies a selection to statistical proteomes to classify protein variants according to some biochemical property, such as the ability to fold correctly into a soluble form or to interact with another protein. Third, Miro uses mass spectrometry (MS) to quantify protein variants and assess differences in the biochemical property for proteins with the ncAA vs. the native amino acid at each position, thereby determining the functional tolerance of the substitution. Because of the global nature of ncAA substitutions, the many biochemical properties that can be assessed, and the capacity of mass spectrometry to quantify peptides, Miro has the potential to identify amino acid substitutions that are likely to be deleterious at an unprecedented scale.

## Results

### Development of the Miro technology

The overall scheme for Miro is shown in Figure 1 for a representative ncAA and biochemical assay. A proline tRNA can be mischarged with a proline analogue, resulting in the random incorporation of the analogue at proline codons and the generation of a statistical proteome (Figure 1a). The statistical proteome is then subjected to a biochemical assay, e.g. a protein affinity purification that separates mistranslated variants of a protein based on their ability to bind to a tagged protein (Figure 1b). Following digestion of the proteins into peptides, mass spectrometry quantifies the abundance of each peptide with a ncAA substitution relative to its native version, before and after affinity purification (Figure 1c). Depletion of a peptide with a ncAA substitution in the purified fraction indicates that the substitution impaired the protein-protein interaction, while enrichment of a substituted peptide indicates that the substitution enhanced the interaction.

**Figure 1.**
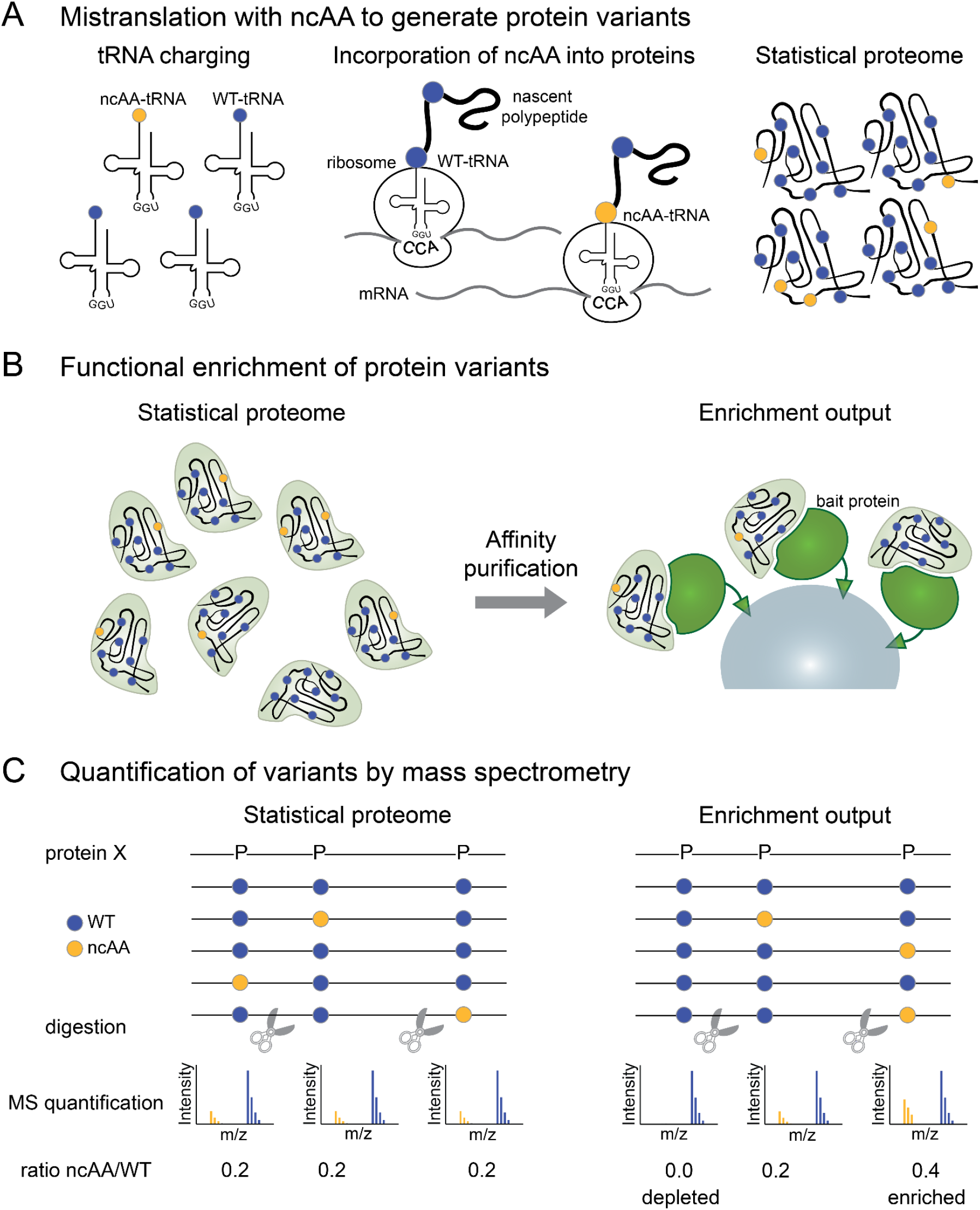
Overview of the Miro method. A) Mistranslation with non-canonical amino acids (ncAAs) to generate protein variants. Non-canonical amino acids are recognized by tRNA synthetases, charged onto their cognate tRNAs, and incorporated randomly into proteins at cognate positions, alongside wild type amino acids, to produce a statistical proteome. B) Functional enrichment of protein variants. The statistical proteome is subjected to a biochemical selection assay to assess the impact of ncAA substitutions on protein function. One example is an affinity purification assay in which protein variants with impaired binding to their partner protein will be depleted from the enrichment output. C) Quantification of variants by mass spectrometry. Mass spectrometry is used to determine the relative abundance of ncAA-containing peptides compared to their wild type cognate versions to determine the incorporation of ncAA at a given position. A comparison of ncAA incorporation before and after the functional enrichment can identify amino acid positions that are functionally important (e.g., in this case for protein-protein interaction). Blue circles represent wild type amino acid proline and yellow circles represent a ncAA analog of proline.

Miro does not rely on modifications to the DNA or to the cellular translation machinery and thus should be generalizable to *in vitro* translation systems and to many organisms and cell types, including bacteria, yeast, and mammalian cells. Because the biochemical assays are carried out on protein lysates, the toxicity of ncAAs towards the cells or translation system should have minimal consequences on the ability to interpret the functional effects of substitutions.

### Incorporation of ncAAs and properties of peptides containing ncAAs

To determine the feasibility of Miro, we first sought to identify ncAAs that both incorporate in the proteome and slow down cell growth, an indication that the ncAA may alter protein function. Additionally, ncAAs with fractional incorporation are ideal, so that individual protein variants contain a small number of substitutions. We screened a panel of 26 ncAAs (Supplementary Table 1) for toxicity and incorporation in *S. cerevisiae* by growing cultures in the presence of a ncAA at one of eight different concentrations and assessing incorporation by mass spectrometry for one or two of these concentrations (Methods). In total, 13 of these ncAAs inhibited yeast growth by at least 25% (Figure 2a, Supplementary Figure 1). Assessing incorporation, 12 ncAAs showed greater than 2% incorporation in the yeast proteome (Figure 2b). Aromatic amino acid analogues with single fluorine substitutions had the highest incorporation, likely due to the conservative nature of the chemical change. We also found several analogues that were toxic without appreciable incorporation, suggesting toxicity mechanisms unrelated to mistranslation. Overall, this screen yielded a set of eight analogues that incorporate at one of seven cognate amino acid positions and are toxic to cells. This set of ncAAs is usable for functional selections in the Miro protocol.

**Figure 2.**
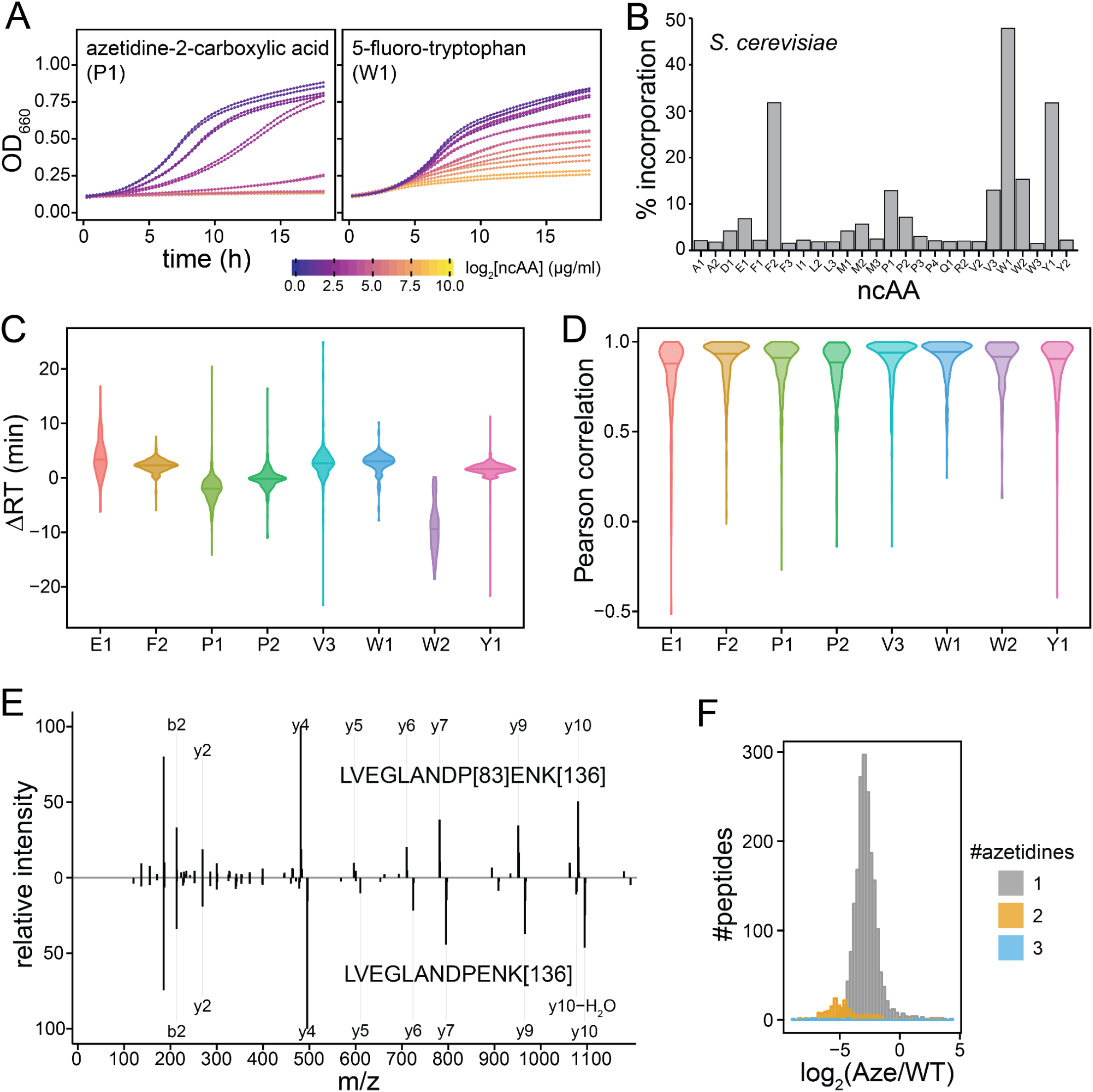
Toxicity and incorporation of ncAA and physicochemical properties of misincorporated peptides. A) *S. cerevisiae* growth curves in the presence of increasing amounts of the ncAAs azetidine (P1) and 5-fluoro-tryptophan (W1) to assess toxicity of non-canonical amino acids (full data in Supplementary Figure 1). B) Incorporation of ncAA on the *S. cerevisiae* proteome. C) Violin plot showing retention time differences between peptides containing ncAAs E1, F2, P1, P2, V3, or W1 and their wild type counterparts. D) Violin plot showing Pearson correlations between MS/MS spectra for peptide precursors containing ncAAs E1, F2, P1, P2, V3, or W1 and their wild type counterparts. E) Mirror MS/MS plot for a charge 2+ precursor of peptide LVEGLANDPENK from Rps12. The spectrum at the top corresponds to the peptide containing azetidine-2-carboxylic acid in place of proline and the bottom spectrum corresponds to the wild type sequence. F) Distribution of P1 incorporation ratios at proline sites on the *S. cerevisiae* proteome. Each data point corresponds to a measured peptide and is colored according to the number of azetidine residues.

Next, we assessed the measurability of peptides containing a ncAA substitution by liquid chromatography coupled to tandem mass spectrometry (LC-MS/MS) by comparing their elution profile and MS/MS fragmentation spectra to those for their matching wild type sequences in LC-MS/MS runs of yeast mistranslated proteomes. Most ncAAs had minimal effects on peptide chromatographic retention time (Figure 2c), and the observed effects were generally in line with the chemical modification introduced by the ncAA. For example, replacement of hydrogen atoms with fluorine tended to increase peptide hydrophobicity and lead to delayed elution (Figure 2c analogues F2, W1, Y1), while the addition of hydroxyl groups had the opposite effect (Figure 2c analogue W2). Furthermore, replacing proline with the 4-membered ring analogue azetidine-2-carboxylic acid (azetidine) advanced elution, due to both decreased hydrophobicity as a result of ring contraction and to a higher propensity towards the cis configuration of the peptide bond (Figure 2c analogue P1, Supplementary Figure 2a-b)^10,11^. The retention time shift of this substitution was dependent on peptide length, relative sequence position, and neighboring amino acid (Supplementary Figure 2c-e).

Peptides containing ncAAs can be identified and the precise position of the substitution localized within the peptide sequence using a similar analysis strategy to that for identifying post-translational modifications, i.e. by allowing a defined mass shift from the protein sequence database on the precursor and some of the fragment ions. This strategy allows us to assess ncAA substitutions at a positional resolution of a single amino acid. MS/MS spectra for wild type peptides and peptides containing a single ncAA substitution were similar, as shown by the high spectra correlations (Figure 2d) and direct spectra comparison for a representative proline-containing peptide (Figure 2e). These similarities indicate that peptide pairs can be reliably quantified and compared at both the full MS and the MS/MS levels.

Finally, in order to accurately quantify ncAA incorporation, we implemented a pulsed-labeling strategy in yeast whereby isotopically-heavy lysine was added to growing cultures at the same time as the ncAA of interest. This strategy mitigates experimental variability due to the chronic toxicity of the ncAA and improves estimates of incorporation ratios across the proteome, specifically by labeling proteins that were synthesized after yeast were exposed to the ncAA. To test this approach, we quantified azetidine incorporation in the yeast proteome. Azetidine incorporation was quantified for 2,112 unique peptides containing one or more proline residues across 789 proteins. Observed incorporation ratios were relatively uniform, with a median incorporation of 10.5% and interquartile range between 5.9 and 18.4%, suggesting that incorporation in yeast is stochastic in nature and, as expected, that the lowest incorporation corresponds to peptides with multiple azetidine substitutions (Figure 2f).

Together, we show that eight ncAAs, when individually added to growth media, were toxic and incorporated stochastically into the yeast proteome by mistranslation. Generally, peptide pairs with and without the ncAA showed similar elution and fragmentation in LC-MS/MS analysis. These results demonstrate the feasibility of the mistranslation approach with ncAAs to assess the functional impact of amino acid substitutions at the scale of the whole proteome. Additionally, because mass spectrometry measurements on Miro are carried out at the peptide level, the impact of substitutions can be assessed at the site level, with the peptides containing a ncAA collectively reporting for all the protein variants containing the ncAA at that site, whereas its wild type peptide counterpart reports for all the variants that do not have the ncAA at that site.

### Effects of substitutions on protein function

We next decided to explore the impact on aspects of protein biology of azetidine mistranslation, given its reported proteotoxic effects^12^, efficient incorporation into the yeast proteome (Figure 2b), and ability to alter protein secondary and tertiary structure^11,13^. We applied Miro with azetidine to four functional readouts: i) the ability of an *in vitro* translated protein to fold and interact; ii) the ability of yeast ribosomal proteins to assemble into the ribosome; iii) the post-translational modification of yeast proteins; and iv) the thermal stability of the yeast proteome.

### Effects of azetidine substitutions on protein structure and interactions

For our initial assay, we applied azetidine to a mammalian *in vitro* translation system synthesizing a protein construct composed of three binding modules: a glutathione S-transferase (GST) domain, a human influenza hemagglutinin (HA) tag, and a polyhistidine (His) tag (Figure 3a). We reasoned that this model system would allow us to maximize proline site coverage and assess overall protein folding and interactions in an affinity purification assay, while largely eliminating potential bias due to degradation or toxicity.

**Figure 3.**
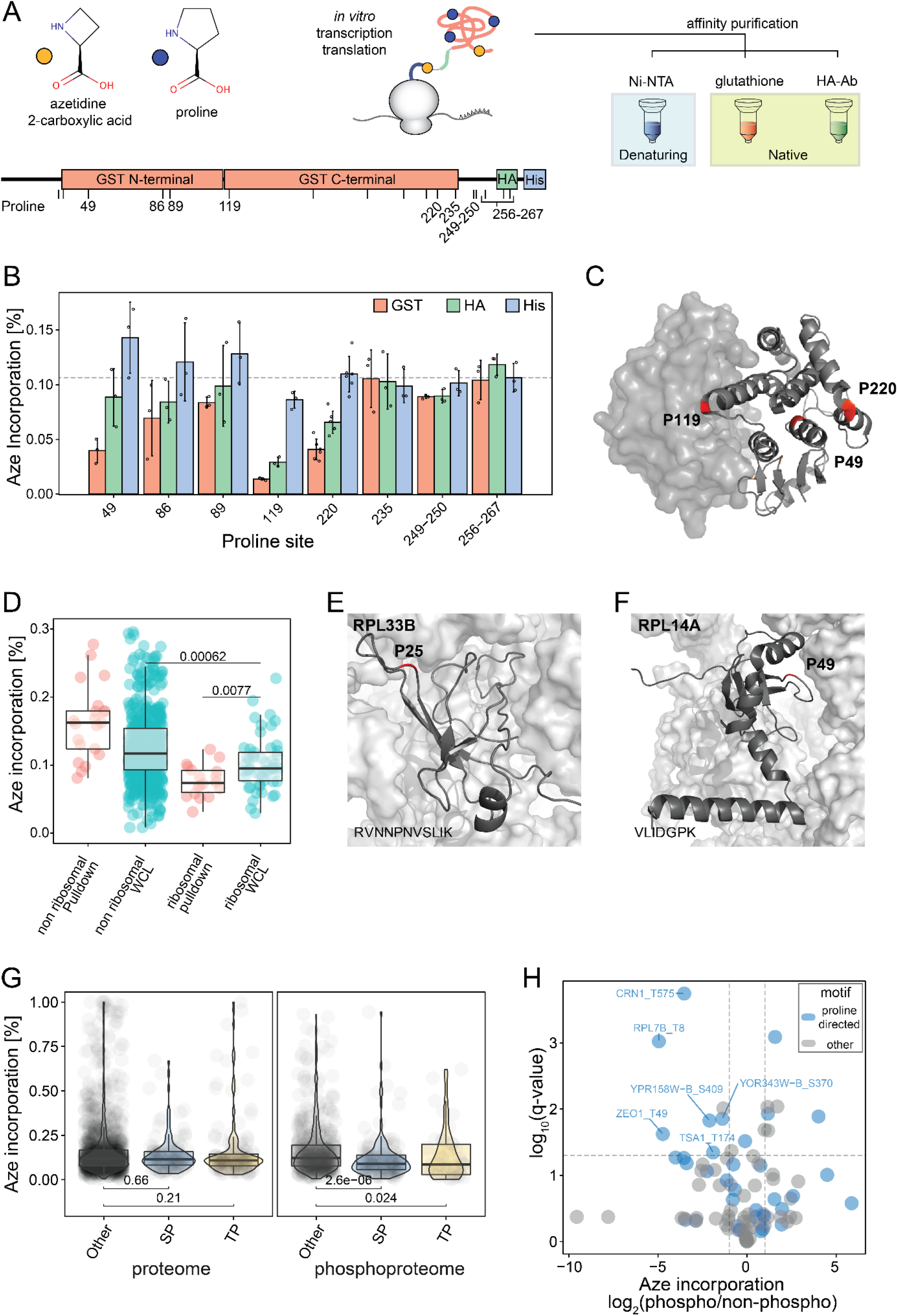
Functional impact of azetidine-2-carboxylic acid incorporation. A) Orthogonal affinity purifications of an HA- and His-tagged glutathione-S-transferase construct that has been mistranslated with azetidine (P1) in a cell-free mammalian in vitro transcription and translation system. B) Relative incorporation of azetidine at proline sites for the three purifications indicated in panel A. C) Surface and cartoon representation of a GST dimer (PDB:1Y6E) showing proline sites with the highest depletion of azetidine in red. D) Boxplot showing incorporation of azetidine at proline sites for the whole yeast cell lysate (WCL) and for the Rpl16B pulldown eluate (PD). Median incorporation rates across replicates per site are shown, and peptides are grouped by fraction and protein of origin (ribosomal or non-ribosomal). The total number of quantified sites per group was: non-ribosomal_PD n=29, non-ribosomal_WCL n=563, ribosomal_PD n=20, ribosomal_WCL n=55 and p-values for group comparisons are computed using a two-sided Wilcoxon test. E, F) Structural context of two proline sites where azetidine incorporation appears as significantly depleted upon ribosome purification: Rpl33B Pro25 (E) and Rpl14A Pro49 (F). The ribosome is shown as surface representation, ribosomal proteins of interest as cartoons (PDB: 4V6I), and relevant proline sites in red. G) Boxplot showing azetidine incorporation into proline positions for the yeast proteome and phosphoproteome. Median azetidine incorporation across replicates is displayed, and peptides are grouped according to the presence of SP, TP or other sequence motifs. Total number of sites quantified for proteome are: (SP motif: n=118, TP motif: n=134, Other: n =1426) and phosphoproteome: (SP motif: n=141, TP motif: n=41, Other: n=488) respectively; and p-values for group comparisons are computed using a two-sided Wilcoxon test. H) Volcano plot displaying azetidine incorporation at proline sites in phosphopeptides relative to their non-phosphorylated counterparts. A t-test was applied to azetidine incorporation rates across replicates for phosphorylated peptides and their non-phosphorylated counterparts (n=40), and p-values were adjusted for multiple hypothesis testing using the Benjamini-Hochberg method.

*In vitro* translation was carried out in the presence of an azetidine excess so as to compete with the proline present in the reaction mixture. We carried out three distinct affinity purifications: a denaturing purification against the histidine tag to assess azetidine incorporation at proline positions throughout the protein, and native affinity purifications against GST and against the HA tag to assess the impact of proline-to-azetidine replacement on the ability of the protein to fold into a native structure (both native purifications) and to interact with its ligand glutathione (only the GST affinity purification) (Figure 3a). Using targeted mass spectrometry, we obtained quantitative information for 11 of the 17 proline sites in the sequence, and calculated intensity ratios between the azetidine-containing peptides and their corresponding wild type forms across all purifications (Figure 3b).

Azetidine incorporation ratios observed in the denaturing purification were uniform across proline positions at ~10% (Figure 3b, Supplementary Dataset 1), validating that ncAA misincorporation in an *in vitro* translation system is largely unbiased. Native purifications, on the other hand, featured several proline positions that were significantly depleted of azetidine, particularly positions located within the GST domains (Figure 3b). This depletion is indicative of azetidine-induced misfolding, which can lead to insolubility prior to an affinity purification. Even though the two native purifications shared similar trends, azetidine depletion generally seemed more pronounced in the GST affinity purification (Figure 3b), with a few site-level differences where azetidine depletion was even more pronounced in the GST purification (Figure 3b).

Thus, the substantial depletion of azetidine at proline 119 in our construct for both purifications is likely the result of its location at the start of the α-helix 4 in the GST C-terminal subdomain. The α-helices 4 and 5 participate in a hydrophobic lock and key arrangement with the other GST monomer^14^, driving the dimerization and crucial to the overall fold (Figure 3c). Conversely, the more extreme depletion of azetidine at proline 49 in the GST purification is likely due to the fact this residue plays an important role in scaffolding the glutathione-binding interface (Figure 3c). Indeed, proline 49 in our construct is located at the N-terminus of α-helix 1 opposite to β-sheet 1 and the conserved catalytic tyrosine residue and mutations at these sites can severely affect glutathione binding^15^. Prolines 86 and 89 are also located in the glutathione binding interface; they were, however, not very sensitive to azetidine substitution. Interestingly, the least affected position was proline 89, perhaps due to the fact that its peptide bond adopts a cis conformation.

### Effects of azetidine substitutions on protein interactions

We next asked whether Miro could identify amino acid positions that are important for the assembly of proteins into a large complex, the ribosome. The yeast cytosolic ribosome is composed of 79 different proteins that assemble with four ribosomal RNAs into a large and a small subunit. We grew *S. cerevisiae* in the presence of azetidine to obtain 5-10% incorporation across the proteome, lysed the cells in native conditions, and affinity-purified the ribosome using an epitope-tagged version of the Rpl16B protein. We analyzed protein mixtures in the whole cell lysate and the ribosome-purified samples to quantify azetidine at each proline position.

We identified over 1200 proteins across two or more replicates. As expected, the Rpl16B pulldown eluate was highly enriched in ribosomal proteins, which contributed almost half of the signal for these samples. This eluate also captured a large fraction of ribosome-associated proteins, as evidenced by Gene Ontology enrichment analysis (Supplementary Figure 3).

In the whole cell lysate sample, we observed lower azetidine incorporation in ribosomal proteins compared to non-ribosomal proteins (p-value=0.0024) (Figure 3d). Azetidine in ribosomal proteins was further reduced in the purified ribosome fraction (p-value=0.035) (Figure 3d), suggesting that in most cases azetidine disrupts ribosomal protein interactions. We identified several proline sites on proteins of the 60S ribosomal subunit with a two-fold depletion of azetidine after pulldown: Rpl33B Pro25 (BH-adjusted p-value=0.014), Rpl14A Pro49 (BH-adjusted p-value=0.021) and Rpl4A P329 (BH-adjusted p-value=0.014). Rpl33B Pro25 is located on the loop connecting strands β1 and β2, which extensively contacts the 25S rRNA (Figure 3e). Similarly, Rpl14A Pro49 is located in the turn that connects the β3 and β4 strands, which interact with both 25S rRNA and Rpl20A (Figure 3f). Finally, Rpl4A P329 mediates a helix-turn-helix motif at the C-terminal extension region, which has been shown to facilitate ribosomal recruitment^16^.

### Effects of azetidine substitutions on protein post-translational modifications

Amino acid substitutions in a substrate protein can affect the ability of modifying enzymes to recognize and catalyze a post-translational modification. We sought to determine whether Miro could be used to identify positions in which a ncAA substitution altered the phosphorylation levels of the protein. In addition to its prominent role in protein structures by forming β-turns, proline is an important residue for recognition by kinases in the CMGC group. These kinases, which include cyclin-dependent kinases (CDKs) and mitogen-activated protein kinases (MAPKs), require a proline at the +1 position of the phospho-acceptor serine or threonine. Therefore, we reasoned that proline substitutions may impact the phosphorylation of SP and TP short linear motifs. To test the effect of such substitutions, we grew *S. cerevisiae* in the presence of azetidine to obtain 5-10% incorporation at proline positions across the proteome.

We measured the content of azetidine at proline sites for whole cell lysate peptides and immobilized metal affinity chromatography (IMAC)-enriched phosphopeptides. Globally, we found that phosphopeptides with a Pro at position +1 to Ser or Thr showed significant depletion of azetidine (Figure 3g). Prolines in other sequence positions relative to a phosphorylated amino acid were not affected by azetidine substitution (Figure 3g).

For 91 proline-containing peptides, we were able to quantify azetidine incorporation in both their phosphorylated and non-phosphorylated forms (Figure 3h) across multiple replicates. Similar to the global trends, the most significantly depleted azetidine incorporations occurred at phosphorylation sites on [S|T]P proline-directed motifs. These instances map to unstructured and readily accessible protein regions, which suggests that azetidine incorporation alters the cis/trans peptide bond conformation balance and likely affects the recognition by kinases or phosphatases^17^.

### Effects of azetidine on protein stability

Lastly, we assessed whether Miro could measure the site-specific effects of ncAA substitutions across an entire proteome. As a proof of concept, we measured the impact of proline-to-azetidine substitutions on protein thermal stability in the *S. cerevisiae* proteome. To do this, we generated a statistical proteome with 5-10% azetidine incorporation in proline positions and implemented thermal proteome profiling (TPP)^18^, a proteomics assay that measures protein thermal stability of thousands of proteins by mass spectrometry. We adapted TPP to produce and compare site-specific thermal denaturation curves for protein variants with and without ncAA substitutions (Figure 4a).

**Figure 4.**
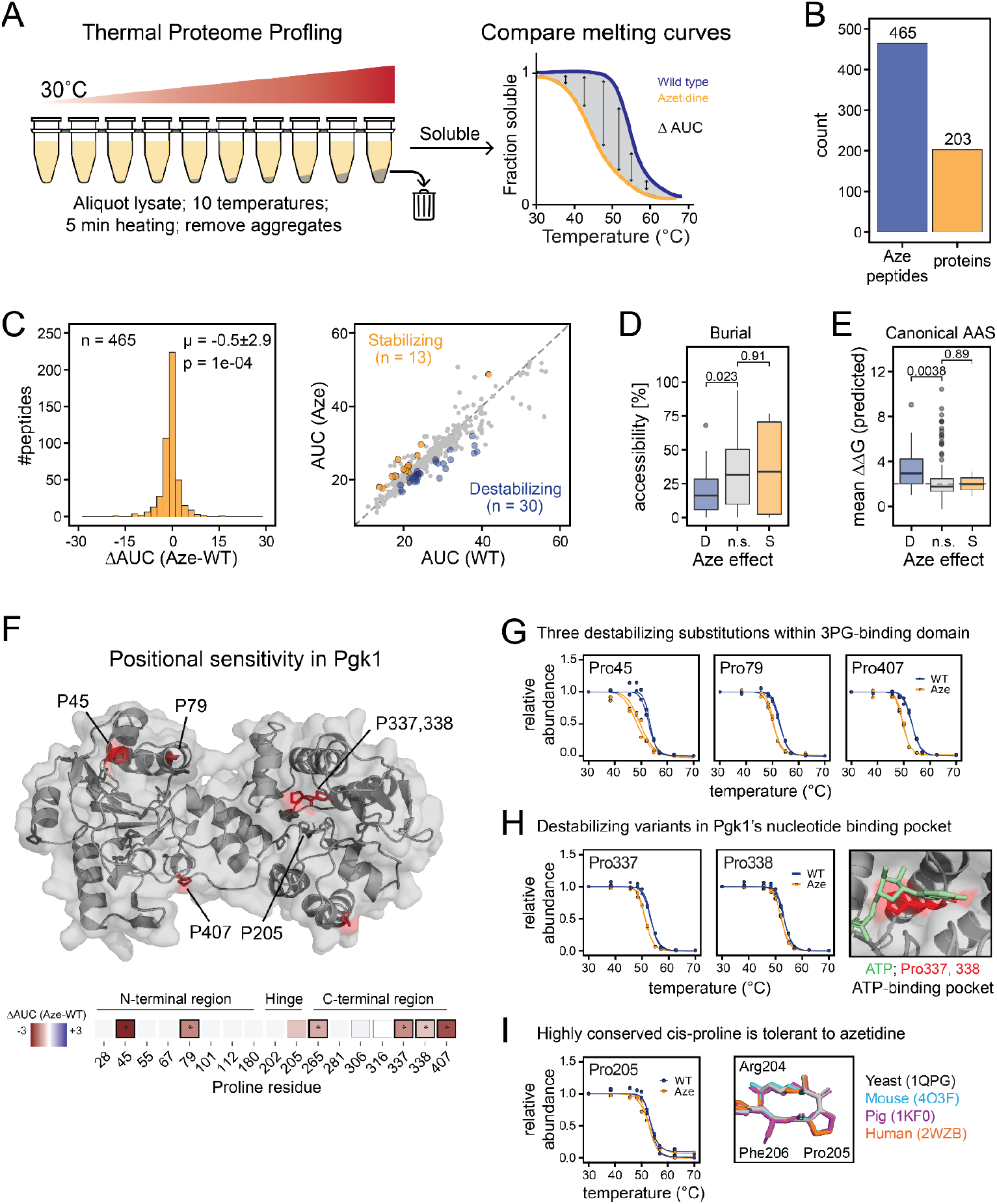
Impact of azetidine substitutions on protein thermal stability. A) Thermal proteome profiling workflow implemented with mistranslated yeast lysates. B) Total number of unique proline-to-azetidine substitutions and total number of proteins with at least one quantified azetidine substitution. C) Left: Distribution of melting curve differences between peptides containing azetidine and matching wild type peptides. Right: Scatterplot showing the relationship between areas under the melting curve (AUC) and the effect of azetidine on protein stability. Substitutions that significantly alter thermal stability (BH-adjusted p-value < 0.01) are colored in yellow (stabilizing) or blue (destabilizing). D) Relationship between the stability effects of proline-to-azetidine substitutions and proline surface accessibility (Destabilizing: n=25; Non-significant: n=364; Stabilizing: n=9). Listed p-values are computed from Wilcoxon rank-sum test. E) Relationship between the stability effects of proline-to-azetidine substitutions and the predicted ΔΔG of natural substitutions at the same prolines (Destabilizing: n=17; Non-significant: n=135; Stabilizing: n=2). Listed p-values are computed from Wilcoxon rank-sum test. F) Structural representation of Pgk1 showing the stability effects of proline-to-azetidine substitutions. Proline residues with an asterisk are statistically significant (BH-adjusted p-value < 0.01). G-I) Melting curves for Pgk1 residues of interest from panel F classified as destabilizing subsitutions in the 3-phosphoglycerate binding domain (G), destabilizing substitutions in the ATP binding pocket (H), and conserved cis-proline (I).

Across two replicates, we identified 1,506 azetidine-containing peptides mapping to 700 yeast proteins, with 660 of these peptides detected and quantified in duplicate in 276 yeast proteins (Supplementary Figure 6a). After stringent filtering (see Methods), we measured changes in stability for 465 unique azetidine substitutions across 203 proteins (Figure 4b). On average, replacing proline with azetidine had a mild destabilizing effect on protein thermal stability (Figure 4c, mean ΔAUC = −0.5 ± 2.9, p-value = 1e-4, two-sided t-test). In total, 43 azetidine substitutions significantly altered melting behavior (permuted non-parametric analysis of response curves^19^, Benjamini-Hochberg corrected p-value < 0.01), most of which were destabilizing (30 destabilizing *vs*. 13 stabilizing) (Figure 4c).

We next analyzed several site-specific structural and evolutionary features for each proline residue where we detected an azetidine substitution in order to understand why azetidine incorporation was destabilizing (D), stabilizing (S), or non-significant (NS). We found that proline residues across all major structural elements (coils, helices, strands, turns) and disordered regions were sensitive to azetidine, although substitutions in β-sheets were generally more destabilizing compared to coiled regions and turns (Supplementary Figure 4c). We also found that destabilizing substitutions occurred at proline residues with lower solvent accessible surface area (Figure 4d, Wilcoxon rank-sum test, p-value = 0.023), and these same sites were predicted to be more sensitive to natural amino acid substitutions (predictions of ΔΔG from mutfunc database^20^; Wilcoxon rank-sum test, p-value = 0.0038, Figure 4e), suggesting that these proline residues may be intolerant to most amino acid changes. Lastly, an unexpected finding was that azetidine sensitivity was not correlated with the evolutionary conservation of proline residues (Supplementary Figure 4c). Taken together, these data suggest that the effect of azetidine may depend less on broad structural features shared across the proteome, and more on local structural and functional contexts specific to each protein.

Next, we asked whether we could identify functional amino acid residues by mapping the effects of azetidine substitutions back to protein structure. We focused on the essential glycolytic enzyme phosphoglycerate kinase-1 (Pgk1), due to its high coverage in our dataset (9 of the 17 possible proline-to-azetidine substitutions) and range of azetidine effects (mean ΔAUC = −1.64 ± 1.40; Figure 4f). Most substitutions had an effect on Pgk1’s stability (6 out of 9 azetidine substitutions significantly destabilizing Pgk1; Figure 4f). Mapping these positions back to Pgk1’s structure reveals slight differences among Pgk1’s domains. Three of the most destabilizing substitutions (azetidine incorporation at Pro45, Pro79, or Pro407) fall in and around the 3-phosphoglycerate-binding domain (Figure 4g). Conversely, the nucleotide-binding domain contains a mix of destabilizing and neutral substitutions, with two destabilizing substitutions falling directly in Pgk1’s nucleotide-binding pocket (Pro337, Pro338) (Figure 4h). Pro205, which has been functionally implicated in Pgk1’s catalytic cycle^21^, was unexpectedly tolerant to azetidine (Figure 4i). Interestingly, this proline adopts a cis conformation in several Pgk1 crystal structures from different species (Figure 4i). Additionally, replacing Pro205 with histidine or phenylalanine, which locks the trans-conformation, has been shown to decrease Pgk1 stability^21^. This result suggests that Pro205 may be tolerant to proline-like analogues that maintain the cis conformation, such as azetidine, and more generally that other cis-conforming proline residues across the proteome may tolerate these types of substitutions (Supplementary Figure 4c).

We also show that Miro can help pinpoint structural elements or regions within proteins that play an important role in protein stability. For example, we found a cluster of destabilizing proline-to-azetidine substitutions within the glycolytic enzyme phosphoglycerate mutase-1 (Gpm1) that all fell within a stretch of four consecutive proline residues (Supplementary Figure 5a). These four residues are all in trans conformation, forming a polyproline-II helix (Supplementary Figure 5a). Azetidine incorporation into polyproline peptides has been shown to influence the cis/trans conformation of the entire helix^11^. While this stretch of prolines has not been previously implicated in Gpm1 stability or function, our data suggest this conformationally-rigid polyproline region is important for Gpm1 stability, and any transition towards a polyproline-I helix (all cis) may be detrimental.

Taken together, these data illustrate how patterns of stability-altering ncAA substitutions within a protein reveal residues and regions important for structure and function. Increasing the depth of substitutions covered in these assays and mistranslating with a variety of ncAAs will bring us closer to interpretable positional sensitivity maps for each protein in the proteome.

## Discussion

Miro is a technology that harnesses mistranslation to produce protein variants *en masse* and mass spectrometry readouts to functionally annotate the consequences of amino acid substitutions on protein function. Using mistranslation with the ncAA azetidine-2-carboxylic acid as a proof of concept, we demonstrate that Miro can identify residues important for protein folding, interaction, and stability.

Mistranslation with ncAAs does not require genetic manipulation. We report a set of ncAAs that incorporate into proteins and can be used in Miro to identify positions that are highly sensitive to substitution. Some amino acids allow for the incorporation of a variety of ncAAs with diverse chemical changes, which should help to interpret the functional consequences of substitutions. For instance, experiments exploring multiple proline analogues with different cis/trans propensities can illuminate protein positions that are sensitive to the conformational context versus general sensitivity to any amino acid substitution.

Miro should be extensible to natural amino acid substitutions by altering the cellular translation machinery to either enhance naturally-occurring mistranslation^22^ or to produce tailored substitutions, for example, by the use of engineered tRNAs^23,24^. Additionally, the same high throughput mass spectrometry assays developed and implemented in Miro could be used to characterize and functionally annotate variant libraries generated for deep mutational scanning experiments^6^.

Similar to other discovery-type mass spectrometry proteomics approaches, Miro does not achieve complete sequence coverage due to sample complexity and the wide range of peptide abundances. In the case of Miro, this limitation is further amplified by the low incorporation of ncAAs necessary to facilitate data interpretation. However, sample enrichment strategies, targeted mass spectrometry acquisition approaches, and future instrumentation improvements should help in the detection of substitution sites and alleviate the coverage issue.

Miro is cost effective, scalable to most types of amino acids, and versatile with respect to the model system in which it can be employed. In this work, we have shown application in yeast and *in vitro* translation reactions; however, we have also screened a subset of these amino acid analogs in *E. coli* and observed similar rates of toxicity and proteome-wide mistranslation. We expect the method to also be adaptable to cultured mammalian cells.

Lastly, Miro is versatile in terms of the protein biochemical properties and functions that can be probed. We have assessed the effects of azetidine substitutions on protein folding, interactions, modifications, and thermal stability, but Miro can readily be extended to assay protein aggregation, enzymatic activity, small molecule binding, and subcellular localization. Further development of Miro with an expanded suite of non-canonical amino acids and the use of engineered tRNAs to carry out natural substitutions will enable the generation of proteome-wide mutational sensitivity maps for human proteins. As companions to genome sequencing efforts, these maps should allow us to interpret the clinical significance of millions of mutations in the human genome. Furthermore, from the standpoint of protein science, Miro will underpin efforts towards detailed sequence-function relationships, enabling the rational design of proteins, including those with enhanced properties for pharmaceutical and biotechnological applications.

## Methods

### Amino acid analogue toxicity screen and incorporation tests

*S. cerevisiae* strain BY4741 was grown overnight in minimal media supplemented with +Ura, +Leu, +Met, +His, 2% glucose at 30°C, diluted into fresh media to a final OD of 0.1, and grown for two additional doublings. At OD of 0.4, cultures were diluted 1:1 in a 96-well plate containing fresh minimal media and a ncAA of interest at one of eight concentrations. The ncAA concentrations used were 2-fold serial dilutions from a starting concentration of 1000 µg/ml (F1, L3, M2, M3), 500 µg/ml (A1, A2, F2, F3, I1, L2, M1, P2, P3, P4, Q1, V2, V3, W2, W3), or 250 µg/ml (D1, E1, P1, R2, W1, Y1, Y2). Optical density of cultures was monitored for 18 h in a temperature-controlled plate reader (BioTek) measuring OD660 every 15 min.

To assess analogue toxicity, we measured the area under the growth curve (AUC) and established relative toxicity to a control growth curve present in each plate. To do this, we first took the average OD_660_ readings from each time point collected for growth curves. We then scaled growth curves between different plates by subtracting the very first OD reading for each growth curve. We then calculated AUC by summing up all the OD readings for each individual condition (i.e. analog, concentration). To assess the effect of analogue treatment on relative growth during this time window, we then divided the AUC values for each condition by the AUC for the control, untreated condition.

Collected growth curves were then used to select ncAA concentrations for incorporation analysis. For ncAAs that showed dose-dependent changes in cell growth, we qualitatively selected an analogue concentration around the IC_50_. For ncAAs that were non-toxic, we chose either 500 µg/mL or 1000 µg/mL. In order to assess incorporation, yeast were grown in 20 ml of minimal media (+Ura, +Leu, +Met, 2% glucose) at 30°C to OD 0.2 at which point both heavy lysine and a ncAA were added to the culture. One exception were cultures exposed to P1, where we assessed incorporation in synthetic complete media (2% glucose) to guide downstream experiments with this analog. Cultures were harvested at OD 1.0 by centrifuging cultures at 2,850 x g and 4°C for 10 min. Cell pellets were resuspended in ice-cold sterile water, pelleted at 10,000 x g and 4°C for 10 min, snap-frozen in liquid nitrogen, and stored at −80°C until cell lysis and sample preparation. One biological replicate was collected for each analog tested. For each analog, we estimate percent incorporation by dividing the number of ncAA-containing peptide-spectrum matches (PSMs) identified in a run and normalizing by the total number of PSMs that could have containing the ncAA substitution (i.e. number of ncAA-containing PSMs plus number of wild-type PSMs containing the cognate amino acid).

### Yeast growth with pulsed heavy lysine and a proline analogue

Yeast strain BY4741 was grown in lysine drop out synthetic complete media (SC-Lys) at 30°C to OD_600_ 0.2, at which point U-^13^C,^15^N-lysine (heavy lysine, K8) proline analogue azetidine-2-carboxylic acid was added at 90-100 µg/ml. Cells were harvested at OD_600_ 1.0 by centrifuging cultures at 2,850 x g and 4°C and decanting growth media. The pellets were resuspended in ice-cold sterile water and pelleted again at 10,600 x g at 4°C. Cell pellets were snap-frozen in liquid nitrogen and stored at −80°C until cell lysis.

### Azetidine-2-carboxylic acid incorporation into in vitro translated protein and pulldown

Mammalian in vitro translation system (Thermo Fisher Scientific IVT Kit #88330) was used to synthesize GST-HA-His protein as described by the manufacturer. Briefly, HeLa cell lysate, reaction mix and accessory proteins were supplemented with pT7CFE1-CGST-HA-His vector, 1mM L-lysine, 1mM L-arginine, protease inhibitors (complete mini EDTA-free, Roche), and 16 mM azetidine-2-carboxylic acid. Reaction proceeded for 8h at 30°C.

In vitro translated protein was purified over 20 µL of anti-HA agarose beads (Sigma-Aldrich), glutathione agarose beads (GoldBio), or 50 µL of Co^2+^-NTA beads (GoldBio). Binding was done in 50 mM Tris buffer pH 8.2, 120 mM NaCl, and elution with 100 mM glycine pH 2.5 and 20 mM glutathione, respectively. Co^2+^-NTA purification was carried out in 50 mM Tris, pH 8.2, 120 mM NaCl, 8M Urea buffer, and eluted with 300 mM imidazole. Purified protein was reduced with 5 mM DTT for 30 min at 55°C, alkylated with 15 mM iodoacetamide for 30 min at RT, and quenched with 15 mM DTT for 15 min at RT. LysC digestion was carried out at a 1:50 enzyme/protein ratio for 2 h at 37°C and pH 8.9. Digestion was quenched with 1% TFA and peptides were desalted over C18 stage tips prior to LC-MS/MS.

### Cell lysis, protein reduction, alkylation and digestion

Frozen cell pellets were thawed on ice and resuspended in lysis buffer (50 mM Tris pH 8.2, 75 mM NaCl, 8 M urea, 50mM β-glycerophosphate, 1 mM sodium orthovanadate, 10 mM sodium pyrophosphate, 50mM NaF, protease inhibitor (complete mini Roche, 1 tablet/10 ml). Cells were lysed at 4°C by repeated vigorous agitation with 0.5mm zirconia/silica beads in a bead beater (Biospec) using 4 cycles of 60 s with 75 s of rest in between cycles. Lysates were cleared by centrifugation at 211 x g for 2 min to remove beads and at 10,600 x g for 8 min to remove cell debris, both at 4°C. Protein concentration was determined by BCA assay (Pierce, Thermo Fisher Scientific). Proteins were reduced with 5 mM DTT at 55°C for 30 min, alkylated with 15 mM iodoacetamide for 30 min at RT, and quenched with 15 mM DTT for 15 min at RT. Protein extracts were digested with lysyl-endopeptidase, LysC (Wako) in 50 mM Tris pH 8.9 overnight at RT and 1:100 or 1:50 enzyme to protein ratio. All samples were acidified to pH 2 by addition of trifluoroacetic acid (TFA) to 0.5% final concentration to inactivate the digestive enzyme.

### Peptide desalting

Peptides were desalted by reversed-phase solid phase extraction over Sep-Pak tC18 cartridges (Waters) in a vacuum manifold as previously described^25^ or stage tips^26,27^ using Empore C18 material (3M), depending on the peptide scale. Sep-Pak tC18 cartridges were equilibrated with sequential additions of 100% acetonitrile (ACN), 70% ACN with 0.25% acetic acid (AA), 40% ACN with 0.5% AA, and 0.1% TFA. Peptide samples were loaded onto the column and washed with 0.1% TFA and 0.5% AA. Peptide samples were eluted by sequential additions of 750 µl of 40% ACN with 0.5% AA, and 750 µl of 70% ACN with 0.25% AA. For stage tips, peptides were applied to conditioned Empore C18 material, washed with 40 µl 0.1% TFA, and eluted with 40 µl 70% ACN with 0.25% acetic acid (AA). All peptide samples were lyophilized prior mass spectrometry analysis, or phosphopeptide enrichment.

### Phosphopeptide enrichment

Phosphopeptide enrichment was performed by immobilized metal affinity chromatography (IMAC) as previously described^25^. Briefly, aliquots of 1 mg peptide were resuspended in 80% ACN, 0.1% TFA and incubated with IMAC beads at a ratio of 1 μl 5% slurry (in 80% ACN, 0.1% TFA) per 10 μg peptides. Phosphopeptide-bound IMAC beads were washed with 80% ACN, 0.1% TFA. Phosphopeptides were eluted with 70% ACN, 1% NH_4_OH, 29% water. Phosphopeptide eluents were stage-tip desalted and lyophilized as described above.

### Pulldown of yeast ribosomal proteins

The strain expressing tagged YNL069C from the MORF collection^28^ was grown in biological triplicate using lysine drop out synthetic complete media (SC-Lys) overnight at 30°C and raffinose as carbon source. Cells were inoculated at OD 0.1 into SC media containing K8, 90 μg/mL azetidine-2-carboxylic acid concentration and 1% galactose to induce Rpl16B expression. Cells were harvested at OD 0.8 by centrifuging cultures at 2,850 x g at 4°C and decanting growth media. Cell pellets were resuspended in ice-cold sterile water and pelleted again at 10,600 x g at 4°C. Cell pellets were snap-frozen in liquid nitrogen and stored at −80°C until cell lysis.

For each biological replicate, cells were resuspended at 4°C in native lysis buffer consisting of 50 mM Tris, 100 mM NaCl, pH 7.5, 0.05% Tween 20 and protease inhibitors (complete mini 1 tablet/10 mL, Roche). Cell suspensions were lysed by four cycles of bead beating (1 min beating, 1.5 min rest). Lysates were centrifuged at 211g for 2 min to remove beads and at 10,600 x g for 8 min to remove cell debris. Supernatants were applied to 60 μL of conditioned lgG Sepharose 6 Fast Flow beads (GE Healthcare) and incubated at 4°C for 3 h. Beads were washed with 50 mM Tris, 100 mM NaCl, pH 7.5, and the bound proteins were eluted with 100 mM glycine pH 2.5. Proteins in the eluate were precipitated with 20% TCA, and their pellet washed with cold 10% TCA solution, followed by cold acetone, and drying. Protein pellet was resuspended with a buffer containing 50 mM ammonium bicarbonate and 10% ACN, and digested for 4 h with 2 μg of lysyl-endopeptidase (LysC). Digestion was quenched with 0.5% TFA and LysC peptides were desalted on styrene divinylbenzene stage tips (3M) and lyophilized.

### Thermal proteome profiling in yeast cell lysates

Two replicates of pelleted yeast (BY4741) containing azetidine mistranslation were resuspended in 650 µl of native lysis buffer (50 mM HEPES pH 7.5, 75 mM NaCl, 2 mM MgCl_2_, protease inhibitors) and lysed by 4 cycles of bead beating (1 minute each with 1 min rest on ice). Lysates were centrifuged for 10 min at 21,000 x g to remove cell debris. Supernatant was collected and transferred to 2 mL tubes. These lysates (2.25 mg/ml concentration) were then aliquoted into PCR tubes on ice. PCR tubes were incubated on a thermal cycler in two phases: first, a 5-min incubation at 30°C; second, a 5-min incubation at 10 different temperatures (30°C, 38.3°C, 45.6°C, 48.3°C, 50.0°C, 52.0°C, 54.7°C, 57.0°C, 62.6°C, 70.0°C) for 5 min. After temperature treatment, lysates were incubated at room temperature for 5 min. All samples were then centrifuged at 17,000 x g 4°C for 1 h. After centrifugation, 75 µl from each temperature-treated sample was mixed 1:1 with denaturing buffer (9M urea, 10 mM DTT, 50 mM HEPES pH 8.9, 75 mM NaCl) and incubated at 55°C for 30 minutes. All samples were then incubated in the dark with 15 mM iodoacetamide for 30 min to alkylate cysteines and the reaction was quenched with 5 mM DTT for 30 min at RT. Protein concentration was measured on an additional aliquot treated at 30°C using the BCA assay.

For each temperature, 75 µg of reduced and alkylated protein lysate was digested with LysC at a 1:50 enzyme:substrate ratio, shaking overnight at RT. Digestion was stopped by addition of 10% TFA to a final concentration of 1.5% TFA. Precipitate was removed by centrifugation, peptides were cleaned up by solid phase extraction on a µHLB Oasis Plate (Waters), and eluates dried down by vacuum centrifugation. 25 µg of dried peptides were resuspended in 100 mM HEPES buffer pH 8.5, 30% acetonitrile, and were labeled with 100 µg of TMT10plex isobaric label reagent (Thermo Fisher Scientific) for 1h at room temperature. The reaction was quenched by addition of 5% hydroxylamine to a final concentration of 0.5% and 30 min incubation. TMT channels corresponding to the different temperatures were pooled together prior to acidification to pH 3 with hydrochloric acid. Acidified peptides were desalted using Sep-Pak tC18 columns (Waters). To increase coverage in Thermal Proteome Profiling experiments, peptides were stage tip fractionated using a high pH reversed phase stepwise elution approach^29^. Briefly, peptides were acidified with 1% TFA, and loaded over conditioned Empore SDB-RPS C18 material (3M), washed and eluted into 4 fractions using a buffer of 20 mM ammonium hydroxide with stepping 5%, 10%, 20% and 80% acetonitrile. Peptide fractions were acidified with 10% FA and dried down by vacuum centrifugation. Samples were analyzed on an Orbitrap Lumos Tribrid mass spectrometer using an SPS-MS3 TMT method^30^.

### Mass spectrometry

Lyophilized peptide, phosphopeptide, and TMT-labeled peptide samples were resuspended in 3% ACN, 4% formic acid and subjected to liquid chromatography coupled to tandem mass spectrometry (LC-MS/MS). Peptide samples were loaded into a 100 μm ID x 3 cm precolumn packed with Reprosil C18 1.9 μm, 120Å particles (Dr. Maisch). Peptides were eluted over a 100 μm ID x 30 cm analytical column packed with the same material housed in a column heater set to 50°C and separated by gradient elution of 8 to 30% ACN in 0.15% FA over 70 min at 350 nl/min delivered by an Easy1000 nLC system (Thermo Fisher Scientific).

Peptides were online analyzed on a Q-Exactive mass spectrometer (Thermo Fisher Scientific). Mass spectra were collected using a data dependent acquisition method. For each cycle a full MS scan (300-1500 m/z, resolution 70,000, AGC target 3e6) was followed by ten MS/MS scans (isolation width 2.0 Da, 26% normalized collision energy, resolution 17,500, AGC target 5e4) on the top 20 most intense precursor peaks.

TMT labeled peptides were loaded onto a 100 μm ID x 3 cm precolumn packed with Reprosil C18 1.9 μm, 120Å particles (Dr. Maisch). Peptides were eluted over a 100 μm ID x 30 cm analytical column packed with the same material housed in a column heater set to 50°C. Peptides were separated by a 120 min gradient of 8-35% ACN in 0.15% formic acid (gradient was optimized for the high pH reverse phase fractions) delivered at 350 nl/min by a nanoACQUITY UPLC (Waters) and online analyzed on a Orbitrap Fusion Lumos Tribrid mass spectrometer (Thermo Fisher Scientific). Mass spectra were collected using a data dependent SPS-MS3 acquisition method^30^, using 5-sec cycles of one full MS scan on the Orbitrap mass analyzer (500-1200 m/z, resolution 60,000, AGC target 5e5), followed by MS/MS scans on the most intense precursor peaks using CID fragmentation and acquisition in the linear ion trap (isolation width of 0.5 Da, normalized collision energy 30, rapid, AGC target 1e4), each followed by an MS/MS/MS scan from coisolating and co-fragmenting the 10 most intense MS/MS fragments, using HCD fragmentation and acquisition in the Orbitrap for reporter ion quantification. (isolation width of 2.5 Da, normalized collision energy 55, resolution of 50,000, 5e4 AGC, max injection time 86 ms).

Analysis of azetidine incorporation on GST-HA-His protein in vitro translated construct was carried out using both data dependent and targeted acquisition approaches on a Q-Exactive instrument. Peptides were eluted over a similar analytical LC set up as above using a gradient elution of 12 to 40% ACN in 0.15% FA over 60 min at 350 nl/min delivered by an Easy1000 nLC system (Thermo Fisher Scientific). DDA methods were acquired as described above, whereas the targeted PRM method consisted of a full MS scan followed by up to 20 targeted MS/MS scans as defined by a time-scheduled inclusion list that included wild type and azetidine 2 carboxylic acid substituted peptides. MS/MS scan was carried out at 35k resolution, 5e5 AGC target, 100 ms maximum injection time, 2 *m*/*z* isolation window, 27% normalized collision energy, centroid mode.

### Mass spectrometry data analysis

Raw files were converted to the mzXML format, and MS/MS spectra were searched against a target/decoy protein sequence database using Comet (version 2015.01, version 2019.01.02)^31^ to identify peptides or MaxQuant (version 1.6.5.0)^32^. *Saccharomyces cerevisiae* (orf_trans_all.fasta downloaded from the Saccharomyces Genome Database in 2016), or human (Uniprot UP000005640 downloaded 2016/09) protein sequence databases were used and tagged bait proteins were included in the database for the affinity purification experiment samples.

Mass tolerance search parameters were adjusted to acquisition instruments following recommendations by Comet source website, i.e. 20 ppm precursor mass tolerance (Orbitrap), 0.02 Da fragment tolerance for MS/MS acquired on an orbitrap mass analyzer and 1.0005 Da tolerance with 0.4 Da offset for MS/MS acquired on a linear ion trap mass analyzer. LysC was selected as the digestive enzyme with a maximum of 2 missed cleavages, constant carbamidomethylation modification of cysteines (+57.0215 Da) and variable modifications of methionine oxidation (+15.9949 Da) and N-terminal acetylation (+42.0106 Da). Variable modifications were also used to search for the incorporation of non-canonical amino acids. For instance, variable modification of −14.0156 Da on proline residues reported for the substitution of proline with azetidine-2-carboxylic acid. Database searches of phosphopeptide samples included variable phosphorylation modification on serine, threonine and tyrosine (+79.9663 Da). Dynamic SILAC samples were searched with light lysine (K0) and heavy (K8, +8.0142 Da) variable modifications in binary mode. TMT-labeled samples were searched with constant modification (+229.1629 Da) on lysines and peptide N-termini. Search results were filtered with Percolator (Percolator version 3.01)^33^ to 1% false discovery rate at the PSM level. Peptide abundance was determined using in-house quantification software to extract MS1 intensity or TMT reporter ion intensities. Protein groups were assembled using ProteinProphet^34^.

For phosphopeptide samples, phosphosite localization was performed using an in-house implementation of Ascore ^35^ using a fragment mass tolerance of 0.4 Da. Phosphosites with Ascore ≥13 were considered localized (> 95% confidence for localization).

Analysis of targeted acquisition data for in vitro translated GST-HA-His pulldown samples was carried out using Skyline (version 3.6.1). Signal extraction was performed on +2, +3, +4, and +5 precursors and +1, +2 b and y fragment ions. Peptide identifications and chromatographic peak boundaries were refined manually, and precursor MS1 intensities were used to calculate azetidine-2-carboxylic acid incorporation. Peptides containing multiple prolines for which azetidine-2-carboxylic acid substitutions could not be resolved chromatographically (n=5), average incorporation rates were calculated.

### Selection of peptides for melting curve fitting

For fitting melting curves, we first applied several stringent filtering criteria. First, we only considered peptides that were fully cleaved, in order to reduce confounding effects due to digestion biases across the temperature range. Second, we only considered peptides containing a heavy lysine, constraining our analysis to the statistical proteome synthesized after exposure to the ncAA. Lastly, we only considered PSMs where at least 5 of the top 10 most intense fragment ions in the MS2 belonged to the assigned peptide, reducing ratio compression from co-isolated precursors. After filtering, TMT reporter ion intensities were transformed into relative fold-changes by normalizing each channel intensity to the channel containing the 30°C control (channel 126). PSMs were consolidated into unique peptides by taking the median fold-change across all PSMs and charge states for each unique peptide.

### Peptide-level melting curve normalization

To account for differences in the amount of material labelled in each channel, we applied a normalization approach similar to a previous approach^36^. Briefly, we selected a set of proteins with relative fold-changes between 0.5 and 1.5 across the entire temperature range, and with a minimum of 3 unique, heavy-labeled, non-redundant, and non-azetidine peptides. We defined this set of proteins as our “non-melting” proteins and used this protein set for sample loading normalization across the entire dataset. Specifically, relative fold-changes for each protein were calculated by taking the median fold-change from all peptides assigned to that protein. We then calculate correction factors so that the median relative fold-change for each replicate and each temperature were equal to 1. These correction factors were then applied across the entire dataset.

### Significance tests for the effect of azetidine substitutions on thermal protein stability

To identify azetidine substitutions that significantly alter protein thermal stability, we compared the melting curves of azetidine-containing peptides with their matching wild type peptides using non-parametric analysis of response curves (NPARC)^19^. In order to estimate the null distribution for our dataset, we took a modified approach to the original implementation of NPARC. We first calculated F-statistics for each peptide pair (degrees of freedom: df1 = 34, df2 = 3). Specifically, peptides that spanned the same ncAA substitution were first consolidated into a site-level quantification by taking the median fold-change across channels. Then, instead of relying on the theoretical F-distribution to calculate a p-value, we generated a permuted dataset of null F-statistics and used these null values to calculate empirical p-values. Specifically, for each observed peptide pair, we generated 500 permuted melting curve comparisons by sampling (without replacement) the relative fold-changes at each temperature, across both replicates and peptide fold-changes. We then fit null and alternative melting curves in R, calculated F-statistics, and combined the 500 permutations across all 465 peptide pairs to generate a dataset of 232,500 permuted curves, each with an associated F-statistic. Any permuted models that failed to converge were first removed before calculating empirical p-values, leaving us with a final dataset of 230,665 null F-statistics (F_null_). We then calculated empirical p-values for each observed F-statistic (F_observed_) using the following formula:

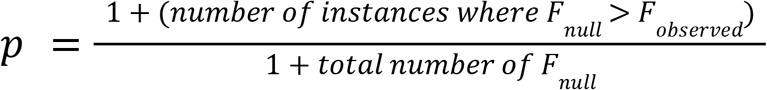

All p-values were corrected for multiple hypothesis testing using the Benjamini-Hochberg method in R.

### Bioinformatics analysis

Gene Ontology Term enrichment for ribosomal pulldown was carried out using YeastMine server (https://yeastmine.yeastgenome.org/). Ontology terms for Biological function, cellular component and molecular function were analyzed for proteins identified in the ribosomal pulldown fraction against a background list of proteins generated from whole cell lysate samples. p-values were corrected using the Holm-Bonferroni method and a 0.05 significance threshold was used.

We assessed the structural and evolutionary context for all proline residues where we detected an azetidine substitution using a variety of computational tools. For all analyses, we only used peptides containing a single azetidine substitution.

For secondary structure analysis, we downloaded PDB files of predicted protein structures from AlphaFold^37^. We assigned secondary structure elements to each residue using DSSP with the Bio3d package in R^38^. The 8-state SSE assignments were then consolidated to 4-state SSE (Helix, Sheet, Turn, Coil) by collectively calling H, G, I states as Helix; E and B states as Sheet; S and T as Turn; and all others as Coil. We calculated relative solvent accessibility by dividing the solvent accessible surface area (SASA) output from DSSP^39^ by a list of maximum SASA per residue^40^. Prediction scores were extracted from each PDB file, and residues with prediction scores less than 70 were excluded from our analysis. Proline residues with omega bond angles between −30° and 30° reside in cis. Disordered predictions for each residue in the yeast proteome were downloaded from http://bioinfadmin.cs.ucl.ac.uk/disodb/. A disordered region was defined as a segment of 30 or more consecutive residues in a protein with a prediction of being disordered.

In order to compare the effects of natural amino acid substitutions with ncAA substitutions, we downloaded the entire *in silico* mutagenesis dataset (both the homology-based and experimental models) from the mutfunc database^20^. We then calculated a mean ΔΔG for each proline residue and used those values as representative of the overall mutational sensitivity of a proline residue.

For our evolutionary analysis, we submitted FASTA sequences obtained from Uniprot (accessed 11 April 2021) to the BLAST server from the command line. Sequences were submitted to BLASTp using default parameters with ‘refseq_protein’ as the search database and ‘Eukarya[ORGN]’ as the entrez query. After searches were complete, we removed any hits with sequence identity less than 30% and E-score > 0.05. We then generated a multiple sequence alignment for each protein in R using the Bio3d and msa package, using the filtered hits as input sequences and default parameters for protein sequence alignments. We then calculated positional entropy within each sequence alignment using the Bio3d package^38^ and the Shannon entropy algorithm. Raw H22 entropy scores were used in our final analysis. We only analyzed positions with more than 100 sequences in the alignment and less than 30% gaps.

## Data availability

The mass spectrometry data have been deposited to the ProteomeXchange Consortium (https://www.ebi.ac.uk/pride/archive/) via the PRIDE partner repository with the dataset identifiers: PXD031230, PXD031233, PXD031255.

## Author contributions

R.A.R.-M. conceived and developed the project, conducted experiments and analyzed data for figures 2 and 3 and associated supplementary figures and supervised the project. K.N.H. conducted experiments and analyzed data for figures 2 and 4 and associated supplementary figures. B.Y.R. conducted ncAA screening experiments and analyzed data. I.R.S., A.S.B, S.Z, Y.Y.L assisted in data analysis. W.S.N. supervised the work of Y.Y.L. and provided funding. S.F. supervised the work of B.Y.R. and S.M.Z. and provided funding. J.V. supervised the work of R.R.A.-M., K.N.H., B.Y.R., I.R.S., and A.S.B. and provided funding. R.A.R.-M. and J.V. supervised the project. R.A.R.-M, K.N.H and J.V. wrote the manuscript and all other authors edited it.

## Acknowledgements

We thank members of the Villén laboratory for enriching scientific discussions. This work is primarily supported by a Medical Research Program grant from the W.M. Keck Foundation (to J.V., S.F., and W.S.N.). This work is supported in part by the University of Washington’s Proteomics Resource (UWPR95794). The work and personnel involved in the project are additionally supported by NIH grants R35GM119536 (J.V.), R01AG056359 (J.V.), and RM1HG010461 (JV. and S.F.); and Human Frontiers Science Program grant RGP0034/2018 (J.V.). K.N.H and I.R.S. were supported by NIH training grant T32HG000035. A.S.B. was supported by NIH training grant T32LM012419. B.Y.R. was supported by NSF Graduate Research Fellowship DGE-1256082. S.M.Z. was a Washington Research Foundation fellow of the Life Sciences Research Foundation.

