## Supplementary for "Proteome-wide identification of amino acid substitutions deleterious for protein function"

### Supplementary Figures

#### Figure S1. Toxicity screen for a library of ncAA

Growth curves for *S.cerevisiae* cultures in the presence of various concentrations of ncAAs (see Supplementary Table 1 for amino acid structures and concentrations).

#### Figure S2. Effects of azetidine-2-carboxylic acid alters cis-trans equilibrium in peptides and alters chromatographic retention time.

A) Histogram of retention time differences between wild type and azetidine-2-carboxylic acid containing peptide. B) Scatterplot showing correlation between chromatographic retention times between s.cerevisiae wild type peptides and their azetidine-2-carboxylic containing counterparts. C) Distribution of retention time differences between wild type and azetidine-2-carboxylic acid containing peptide binned by peptide length. D) Scatter plot of retention time differences between wild type and azetidine-2-carboxylic acid containing peptides and relative position of misincorporated proline site binned and colored by peptide length. E) Boxplot of retention time differences between wild type and azetidine-2-carboxylic acid containing peptide and binned by dipeptide motif where Aze misincorporation occurs.

#### Figure S3. Proteins in the ribosome pulldown.

A)Significant Gene ontology terms enriched in ribosome purification. B) Venn diagram displaying the number and overlap of protein identifications for whole cell lysate and ribosome pulldown eluate fractions, as well as the coverage for the ribosome.

#### Figure S4: Thermal proteome profiling on yeast lysates with azetidine-2-carboxylic acid misincorporation.

A) Summary statistics of identified unique peptides containing azetidine-2-carboxylic acid and corresponding proteins across two replicate thermal proteome profiling experiments. B) The number of singly- and doubly-modified azetidine substitutions detected. C) Associations between structural and evolutionary features with azetidine sensitivity. From left to right: secondary structure (Coil: 162; Turn: 109; Helix: 96; Sheet: 31), structure disorder predictions (Disordered: 41; Ordered: 409), evolutionary conservation measured as raw Shannon entropy values (Destabilizing: 23; Non-significant: 358; Stabilizing: 9), and proline conformation (Cis: 17; Trans: 381). Listed p-values above each comparison are from Wilcoxon-ranked sum test.

#### Figure S5. A conserved polyproline region is important for Gpm1 stability in yeast.

A) Crystal structure (PDB ID: 1BQ4) of tetrameric yeast phosphoglycerate mutase (Gpm1) with the positions of destabilizing substitutions highlighted in red. B) Positional sensitivity map of Gpm1, which includes differences in melting curves ($\Delta$AUC), predicted effects of natural substitutions ($\Delta\Delta$G), and residue conservation across other eukaryotic Gpm1 orthologs. C) Melting curves for two proline-to-azetidine substitutions within a conserved polyproline helix in Gpm1 that decrease protein thermal stability.

### Supplementary Figure 1

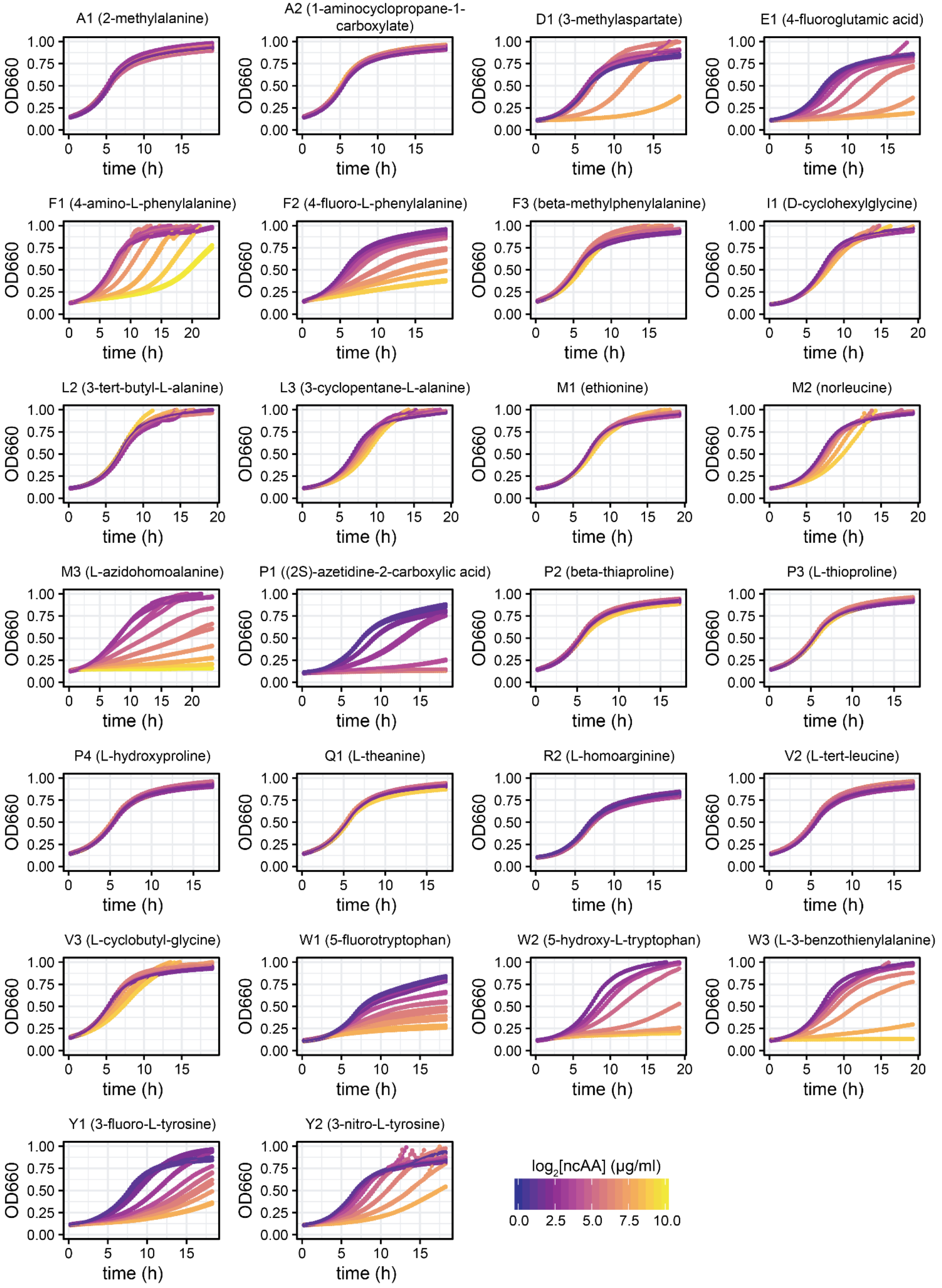

Supplementary Figure 2

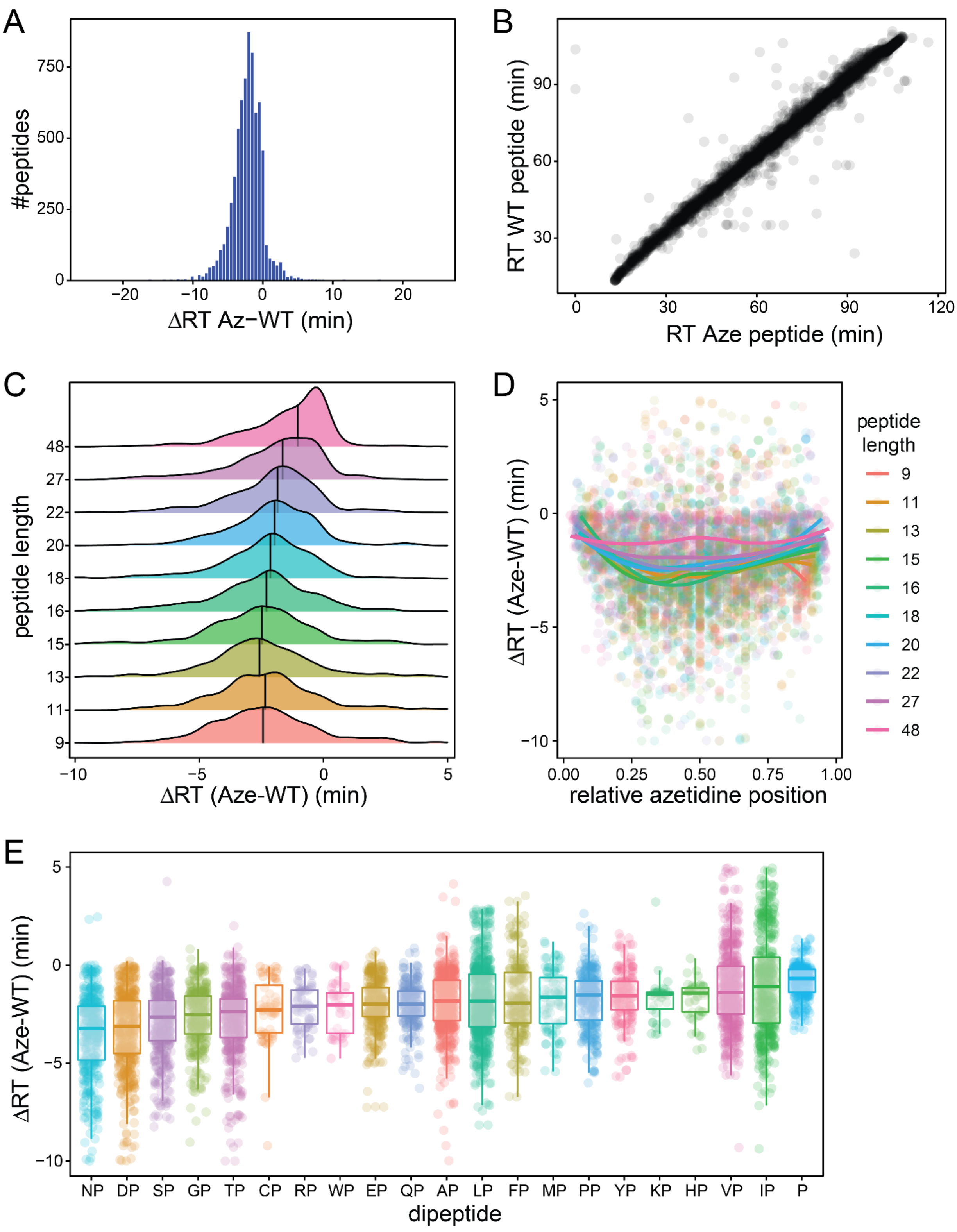

Supplementary Figure 3

A

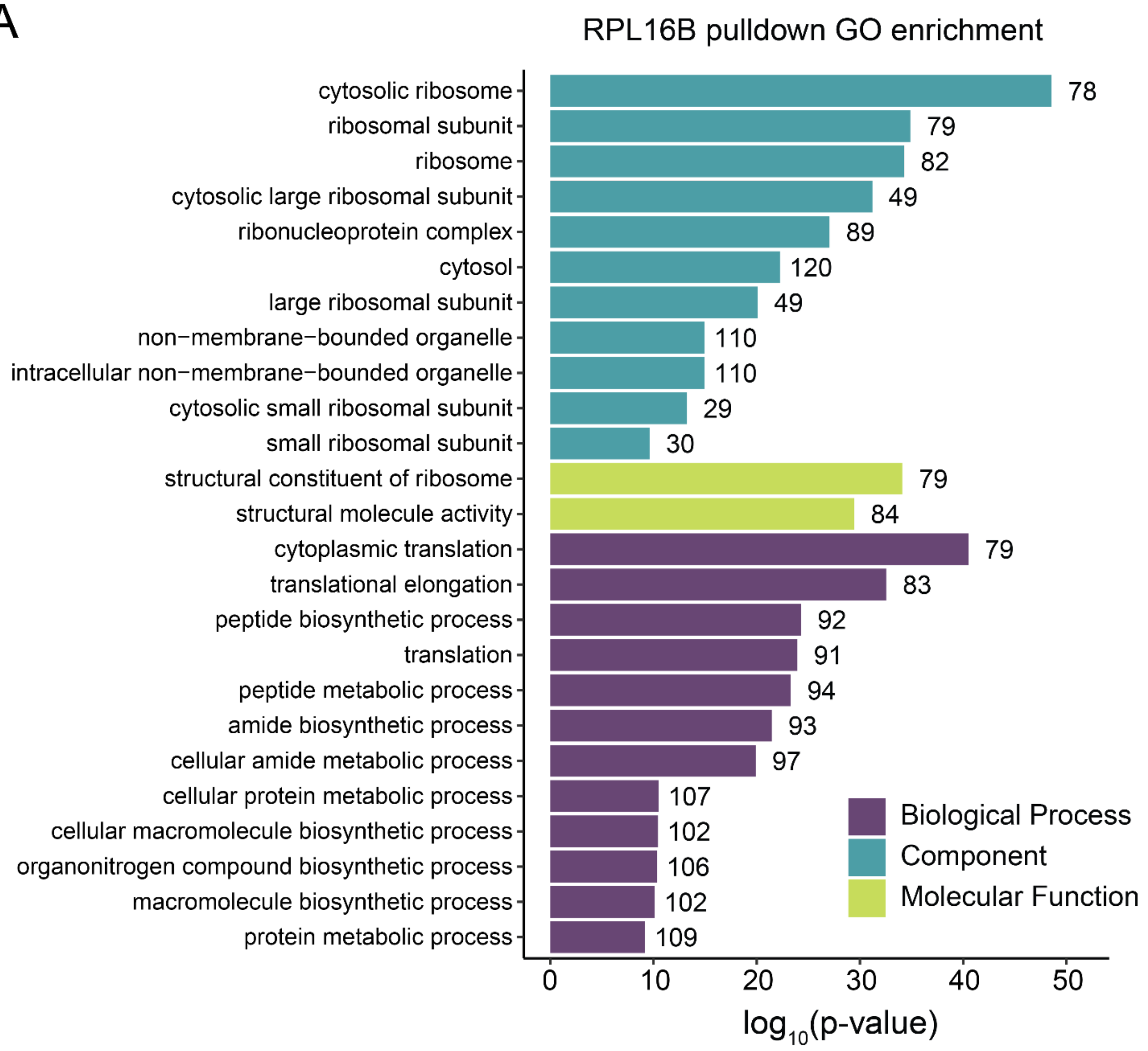


B

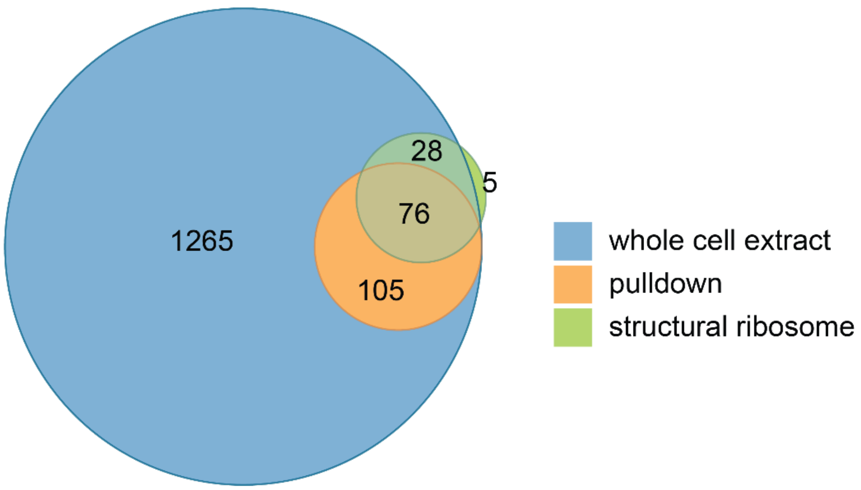

Supplementary Figure 4

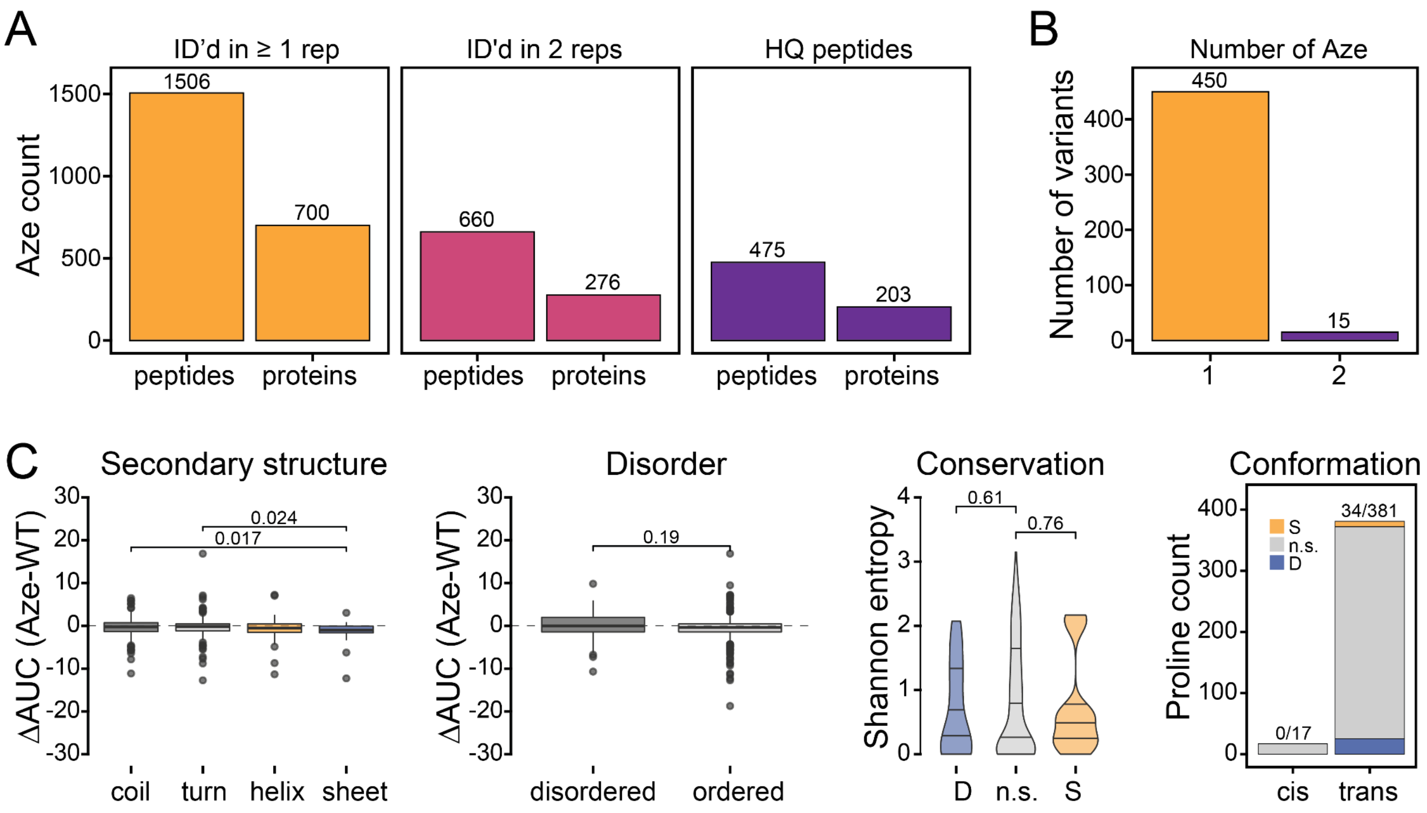


Supplementary Figure 5

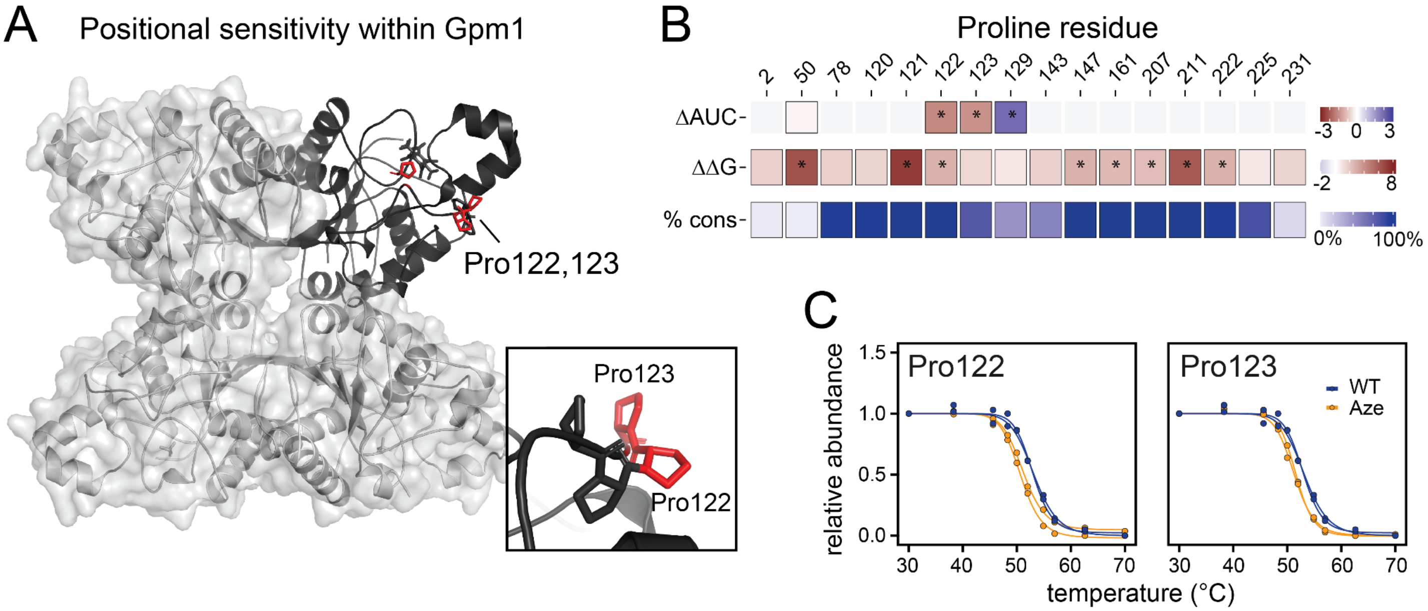

#### Supplementary Tables

Table S1. Library of ncAA with structures and concentrations used in the toxicity and incorporation screens.

| Symbol | Name | Structure | Growth screen concentrations (µg/ml) | Incorporation concentration (µg/ml) |
|---|---|---|---|---|
| A1 | 2-methylalanine | 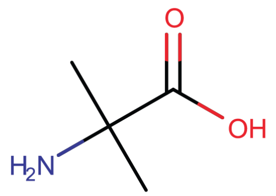 | 3.91<br>7.81<br>15.63<br>31.25<br>62.5<br>125<br>250<br>500 | 500 |
| A2 | 1-aminocyclopropane-1-carboxylic acid | 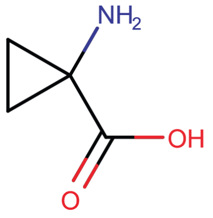 | 3.91<br>7.81<br>15.63<br>31.25<br>62.5<br>125<br>250<br>500 | 500 |
| D1 | 3-methylaspartic acid | 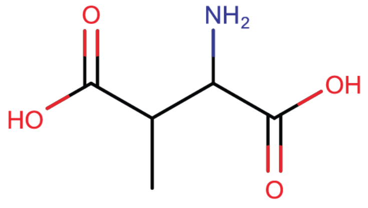 | 1.96<br>3.91<br>7.81<br>15.63<br>31.25<br>62.5<br>125<br>250 | 100 |
| E1 | 4-fluoroglutamic acid | 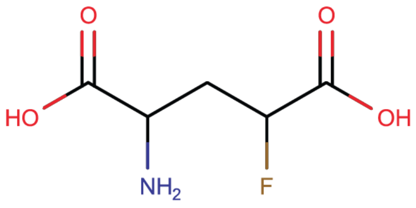 | 1.96<br>3.91<br>7.81<br>15.63<br>31.25<br>62.5<br>125<br>250 | 80 |

| | | | | |
|---|---|---|---|---|
| F1 | 4-amino-L-phenylalanine | 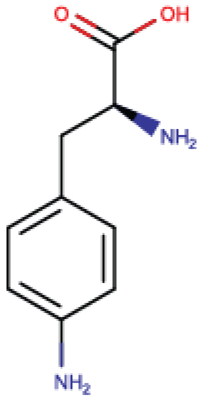 | 7.81<br>15.63<br>31.25<br>62.5<br>125<br>250<br>500<br>1000 | 250 |
| F2 | 4-fluoro-L-phenylalanine | 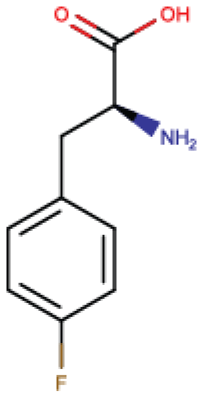 | 3.91<br>7.81<br>15.63<br>31.25<br>62.5<br>125<br>250<br>500 | 30 |
| F3 | beta-methylphenylalanine | 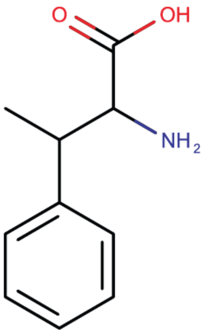 | 3.91<br>7.81<br>15.63<br>31.25<br>62.5<br>125<br>250<br>500 | 500 |
| I1 | D-cyclohexylglycine | 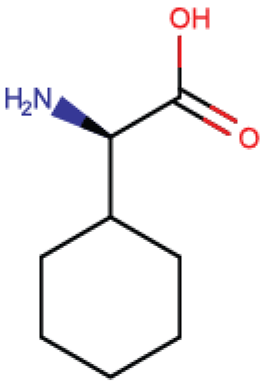 | 3.91<br>7.81<br>15.63<br>31.25<br>62.5<br>125<br>250<br>500 | 1000 |

| | | | | |
|---|---|---|---|---|
| L2 | 3-tert-butyl-L-alanine | 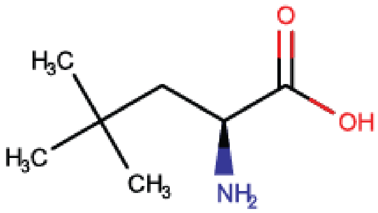 | 3.91<br>7.81<br>15.63<br>31.25<br>62.5<br>125<br>250<br>500 | 1000 |
| L3 | 3-cyclopentane-L-alanine | 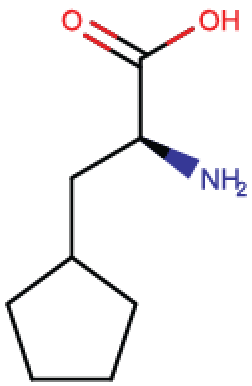 | 7.81<br>15.63<br>31.25<br>62.5<br>125<br>250<br>500<br>1000 | 1000 |
| M1 | DL-ethionine | 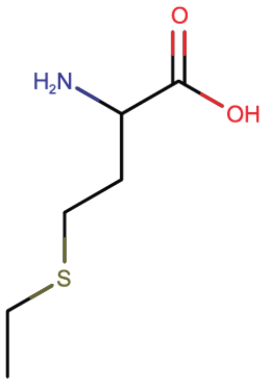 | 3.91<br>7.81<br>15.63<br>31.25<br>62.5<br>125<br>250<br>500 | 1000 |
| M2 | DL-norleucine | 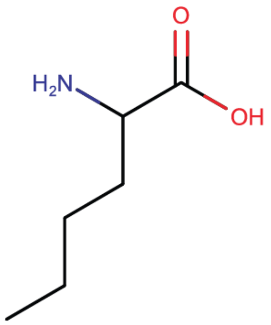 | 7.81<br>15.63<br>31.25<br>62.5<br>125<br>250<br>500<br>1000 | 1000 |

| | | | | |
|---|---|---|---|---|
| M3 | L-azidohomoalanine | 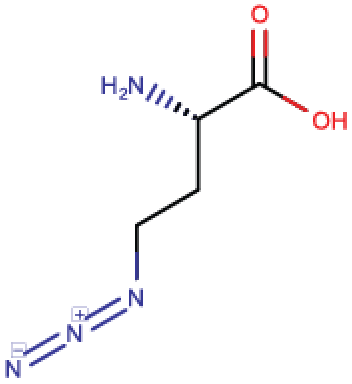  | 7.81<br>15.63<br>31.25<br>62.5<br>125<br>250<br>500<br>1000 | 80 |
| P1 | (2S)-azetidine-2-carboxylic acid | 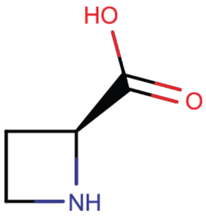  | 1.96<br>3.91<br>7.81<br>15.63<br>31.25<br>62.5<br>125<br>250 | 100 |
| P2 | beta-thiaproline | 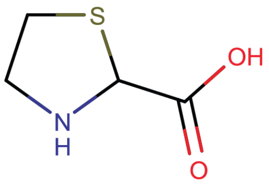  | 3.91<br>7.81<br>15.63<br>31.25<br>62.5<br>125<br>250<br>500 | 500 |
| P3 | L-thioproline | 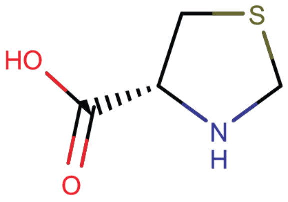  | 3.91<br>7.81<br>15.63<br>31.25<br>62.5<br>125<br>250<br>500 | 500 |
| P4 | L-hydroxyproline | 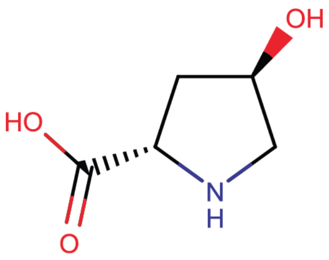  | 3.91<br>7.81<br>15.63<br>31.25<br>62.5<br>125<br>250 | 500 |

| | | | | |
|---|---|---|---|---|
| | | | 500 | |
| Q1 | L-theanine | 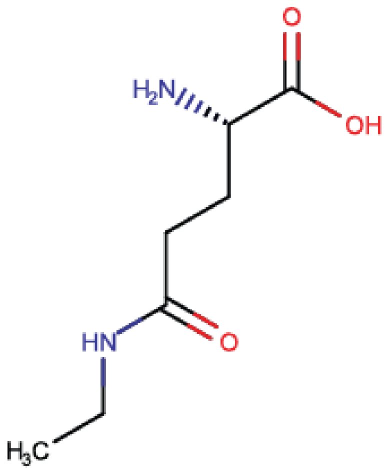 | 3.91<br>7.81<br>15.63<br>31.25<br>62.5<br>125<br>250<br>500 | 500 |
| R2 | L-homoarginine | 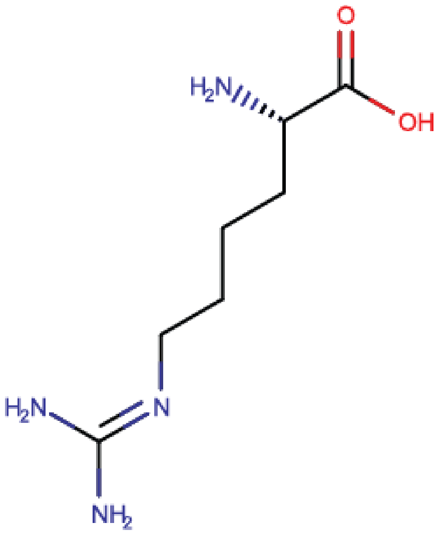 | 1.96<br>3.91<br>7.81<br>15.63<br>31.25<br>62.5<br>125<br>250 | 500 |
| V2 | L-tert-leucine | 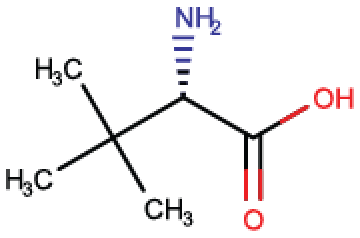 | 3.91<br>7.81<br>15.63<br>31.25<br>62.5<br>125<br>250<br>500 | 500 |

| | | | | |
|---|---|---|---|---|
| V3 | L-cyclobutyl-glycine | 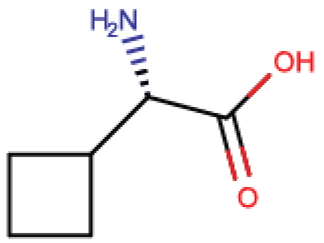 | 3.91<br>7.81<br>15.63<br>31.25<br>62.5<br>125<br>250<br>500 | 250 |
| W1 | 5-fluorotryptophan | 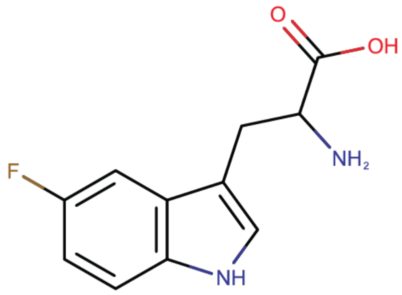 | 1.96<br>3.91<br>7.81<br>15.63<br>31.25<br>62.5<br>125<br>250 | 15 |
| W2 | 5-hydroxy-L-tryptophan | 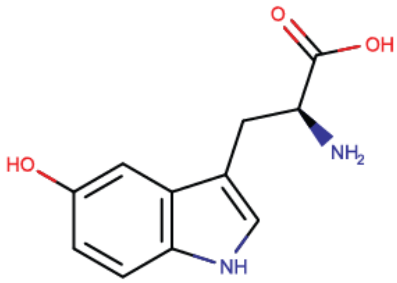 | 3.91<br>7.81<br>15.63<br>31.25<br>62.5<br>125<br>250<br>500 | 30 |
| W3 | L-3-benzothienylalanine | 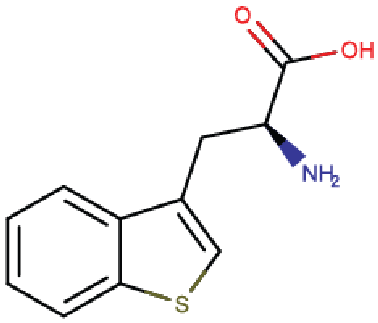 | 3.91<br>7.81<br>15.63<br>31.25<br>62.5<br>125<br>250<br>500 | 80 |

| | | | | |
|---|---|---|---|---|
| Y1 | 3-fluoro-L-tyrosine | 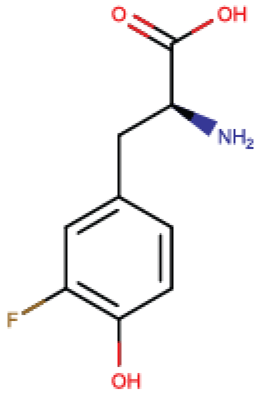 | 1.96<br>3.91<br>7.81<br>15.63<br>31.25<br>62.5<br>125<br>250 | 5 |
| Y2 | 3-nitro-L-tyrosine | 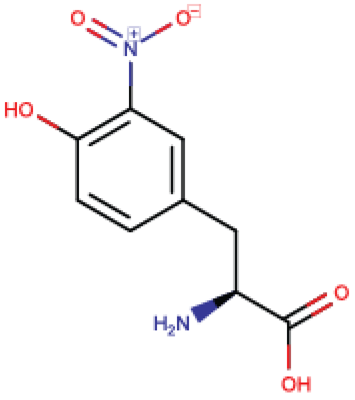 | 1.96<br>3.91<br>7.81<br>15.63<br>31.25<br>62.5<br>125<br>250 | 60 |
